# A Novel Method to Characterize Functional Human Embryonic Stem Cells Derived Cardiomyocytes

**DOI:** 10.1101/2025.10.17.683184

**Authors:** Mohammad Reza Hashemzadeh, Samaneh Aghajanpour, Nasser Aghdami, Reza Aflatoonian

## Abstract

**Background:** As key factors in innate immunity and activators of signals that are critically involved in the commencement of adaptive immune responses, Toll-Like receptors (TLRs) leave a trace in heart diseases, cardiomyocyte (CM) protection, contractility, and maturation. Human embryonic stem cells (hESCs) have the potential to provide an unlimited source of cardiomyocytes, being invaluable resources for cell therapy to restore heart function after damage.

**Objectives:** The main goal of this novel study, TLRs expression during cardiomyocyte differentiation, is to determine whether this assessment can be considered a potential procedure to characterize functional cardiomyocytes.

**Methods:** Royan H5 and Royan H6 hESC lines were cultured in suspension and differentiated into CMs according to the Laflamme protocol, and hESC-CMs were characterized by immunostaining for cytoskeletal proteins. The gene expression for TLR2, TLR3, TLR4, TLR5, and TLR9 was evaluated by RT-PCR and Q-PCR during differentiation. Moreover, the secretion of interleukin (IL)-6 and IL-8 was measured by enzyme-linked immunosorbent assay (ELISA).

**Results:** The cell membrane receptors, TLR2 and TLR4, both represented the downward trend after day 14. However, the expression of endosomal representative receptors (TLR3 and TLR9) surged during differentiation and hit the highest point in fully differentiated CMs.

**Conclusion:** It can be concluded that the generated CMs in this study are functional due to the expression pattern of TLRs being in favor of obtaining desired CMs. Meanwhile, manipulation and optimization of such receptors’ expression is suggested in subsequent studies to obtain highly mature CMs as a gold candidate in cardiac cell-based therapy.

## 1. Introduction

Human pluripotent stem cells like embryonic stem cells (ESCs) and induced pluripotent stem cells (iPSCs) play a pivotal role in studying human diseases due to both self-renewal and to differentiate into any cell types in the body. As such, hESCs can potentially provide an unlimited supply of even highly specialized cells like cardiomyocytes (CM) to restore cardiovascular functions that have been damaged [1–3].

Cardiomyocytes derived from human embryonic stem cells (hESCs) potentially offer large numbers of specialized cells with contraction as a primary function [4]. Increasing evidence demonstrates that CMs can respond to danger signals with a directed and complex inflammatory and functional response [5]. This response involving cytokines, chemokines, and the subsequently recruited leukocytes, as well as cell surface adhesion molecules, leads to decreased cardiomyocyte contractility, thereby modulating peak systolic stress and strain, which may conceivably impact repair processes [6–8].

The first steps that trigger the cardiomyocyte response to danger signals are directed by Toll-like receptors (TLRs). The first steps that trigger the cardiomyocyte response to danger signals are directed by Toll-like receptors (TLRs). TLRs represent a family of pattern recognition receptors that serve to recognize molecular patterns associated with pathogens, induce the activation of several kinases and NF-κB upon binding of their ligands. They play an essential role in innate immunity [9, 10] and induce an innate immune response to tissue damage by responding to endogenous ligands released from damaged tissues or necrotic cells [8]. Even alterations of the extracellular matrix due to tissue destruction can activate the innate immune system through TLR-mediated pathways [11]. Specialized tissues such as skeletal muscle [12], initially thought to be bystanders in the immune response to pathogens, have recently been found to be active participants in response to TLR ligation. TLR2 was found to be increased in the heart tissues of mice with HF under isoproterenol (ISO) challenge [13]. Interaction between TLR4 and fibroblast growth factor 10 (FGF10) has been represented in the transcriptome of human induced pluripotent stem cells-derived cardiac progenitor cells (hiPSC-CPCs) [14].

Myocardial infarction (MI) is the most common and clinically significant form of acute cardiac injury and results in ischemic death of a large number of cardiomyocytes [15]. In addition to reporting an increased level of TLR9 expression in the end-stage heart failure patients after ischemic heart disease, the induction of key TLR9 pathway proteins in a mouse *in vivo* model of myocardial ischemia has been reported [16]. Meanwhile, in progression between coronary atherosclerosis and MI, TLR2 may have a main role and a beneficial screening value for MI [17]. Cells dying by necrosis release their intracellular contents and initiate an intense inflammatory response by activating innate immune mechanisms. According to recent studies, various immune responses occur in different experimental models of cardiac regeneration, and restraining these immune responses may improve cardiac repair [18]. Evidence suggests that TLR-mediated pathways play a significant role in triggering the post-infarction inflammatory response by activating the Nuclear factor (NF)-κB system. For instance, TLR4 plays an important role in myocardial inflammation [19], and modulation of TLR3 can ameliorate heart injury in heart attack [20], which develops and occurs in cardiovascular diseases (CVDs) through the regulation of different cytokines [21]. Generally speaking, emerging data indicate that TLR signaling pathway molecules are involved in the progression of heart failure, so that TLR2, TLR3, TLR4, and TLR9 are activated in the failing heart [22].

Accordingly, the role of TLRs in the cardiovascular system and its related diseases is almost clear. Not only is the evaluation of TLRs expression during CM differentiation a novel study, but considering whether this appraisal can introduce a new way to characterize functional cardiomyocytes was also the goal of our study. Regarding the pivotal role of TLRs in heart diseases, cardiomyocyte contractility, protection, and especially maturation, obtaining functionally differentiated cardiomyocytes characterized by a TLR expression pattern can be a reliable candidate for cell-based therapy after heart disease.

## 2. Methods

### 2.1. Suspension cell culture and differentiation protocol

Suspension culture of hESCs (Royan H5 and Royan H6) was performed according to Larijani *et al*. protocol [23]. Briefly, cells were treated with 10 mM ROCK inhibitor Y-27632 (Sigma-Aldrich, Y0503) 1 h before dissociation from Matrigel. Cells were then washed with Ca ^2+^ - and Mg ^2+^ -free phosphate-buffered saline (PBS; Gibco, 21600-051) and incubated with 0.05% trypsin at 37 °C for 4–5 min. Dissociated cells were transferred into non-adhesive bacterial plates (60 mm; Griner, 628102) at 156104 viable cells/ ml in hESC medium that had been conditioned on mouse embryonic fibroblasts (MEFs) [24], which contained 10 mM ROCK inhibitor. After 2 days, half of the medium was replaced by the hESC medium conditioned on MEFs. The medium was changed every other day. Differentiation of the cells into cardiomyocytes in suspension was performed according to the Laflamme *et al*. protocol [25] with some modifications. Concisely, 6-day-old spheres were treated with 100 ng/ml Activin A for 1 day in RPMI medium (Gibco, 51800035) supplemented with 2% B27 without vitamin A, followed by 4 days of 10 ng/ml BMP4. At day 5, the spheres were plated on gelatin-coated plates in RPMI/B27 medium without cytokines. Beating clusters were observed 5 days post-plating. The recombinant protein LIM homeodomain transcription factor, Islet1 (rISL1), was added from days 1–8 after initiation of differentiation induction. All experiments with hESCs were performed under the supervision of the Institutional Review Board and the Institutional Ethical Committee of our institute.

### 2.2. RT-PCR and gel electrophoresis

Total RNA was isolated from the cells using TRI reagent (MRC, Cincinnati, OH), and contaminating DNA was removed using DNase I treatment. Expression of mRNAs for human TLRs as well as beta-actin was assessed by first-strand cDNA synthesis from 5g of total RNA by extension of oligo (dT) primers with 200 U Moloney murine leukemia virus reverse transcriptase (MMLV-RT; Promega, Madison, WI). The polymerase chain reaction (PCR) program was performed using the prepared cDNA, primers for Toll-like receptors (Table 1) and Platinum Blue PCR Super Mix (Invitrogen) under the following conditions: primary denaturation was 95°C for 5 minutes; then 35 cycles of 95°C for 45 seconds, 60°C for 45 seconds and 72°C for 45 seconds; at the end final extension was 72°C for 30 second. All PCR products were size fractionated by 2% agarose gel electrophoresis, and DNA bands were visualized by ethidium bromide staining. GAPDH gene was used as an internal control.

**Table 1.**
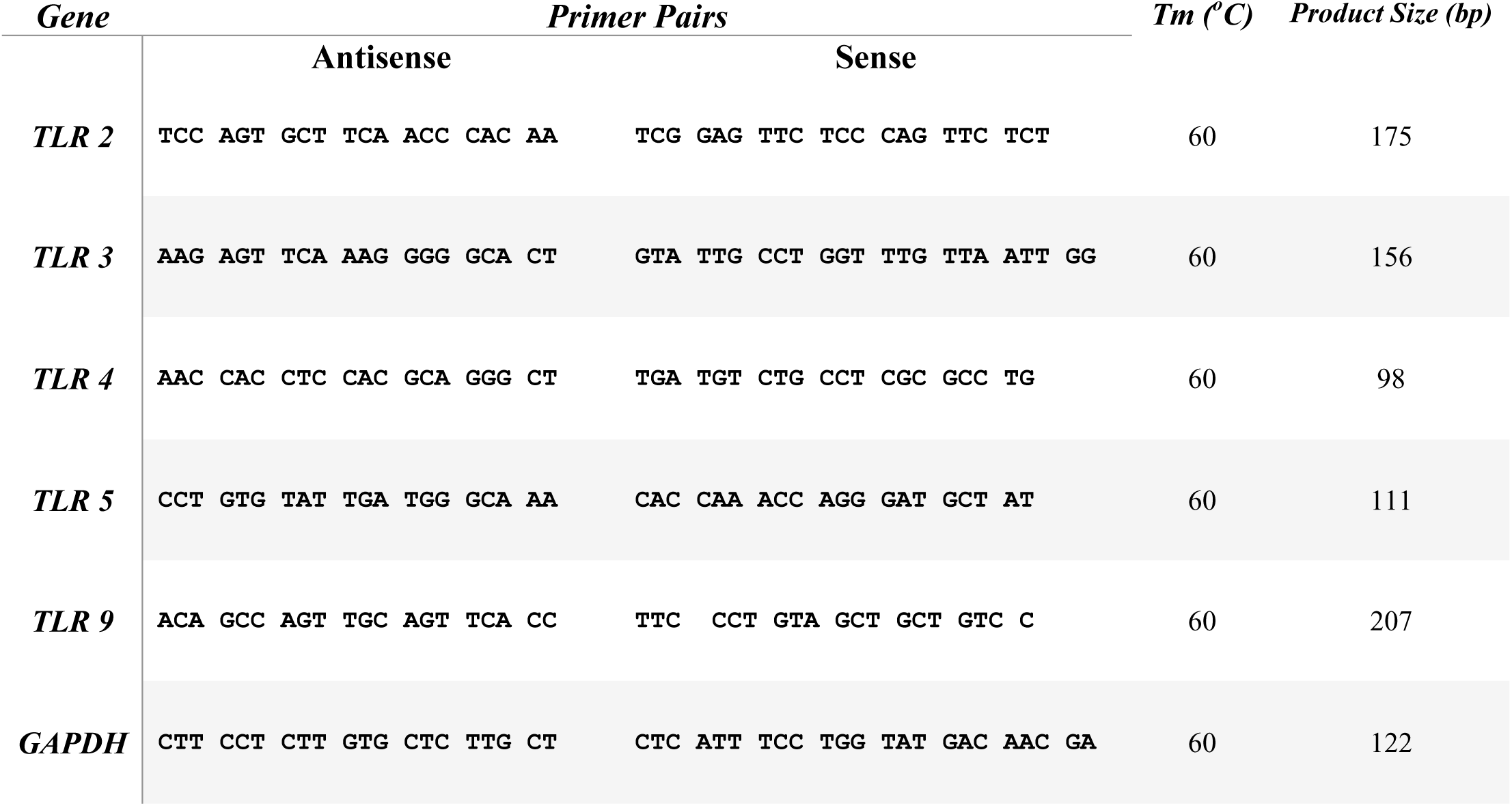
Sequence of Primers.

### 2.3. Quantitative Polymerase Chain Reaction (Q-PCR)

Quantitative real-time PCR was performed using the prepared cDNA, primers for TLRs, and human GAPDH. The forward and reverse primer sequences used are depicted in Table 1. All experiments included negative controls with no cDNA. SYBR Green Jump Start (Sigma) master mix was added to each well of the PCR plate (10 μl of SYBR Green, 7 μl of water, 2 μl of primers, and 1 μl of cDNA), and PCR was performed under the following conditions: 40 cycles at 95°C for 30 s, 60 °C for 30 s, and 72 °C for 30 s. Samples were run in triplicate. Results were analyzed using an iCycler (Bio-Rad Laboratories, Hemel Hempstead, UK).

### 2.4. Immunostaining of ESCs-CM

After collecting of beating spheroids on the 25th day of differentiation, they were washed three times with PBS. By 0.05% trypsin/EDTA and dissociated into single cells in an incubator at 37 °C for 5 min. Then the culture of individualized cells on Matrigel-coated chamber slides in differentiation medium without SMs was done. The attached cells, after 48 h, were washed with PBS, fixed with 4% (w/v) paraformaldehyde at room temperature for 15 min, and washed once with washing buffer (PBS/0.1% Tween 20) respectively. Cell permeabilization was done by 0.2% Triton X-100 in PBS for 15 min, and then blocked with 5% (v/v) bovine serum for 1 h. Primary antibodies were diluted in a ratio 1:100 with blocking buffer included PBS, 5% bovine serum, and 0.2% Triton X-100. Then, diluted antibodies were added to cells and incubated overnight at 4 °C. The next day, the cells washed 3 times with washing buffer for 5 min. interval. Diluted secondary antibodies in blocking buffer (1:500) were then added to cells and incubated for 1 h at RT. Finally, the cells were washed 3 times with washing buffer at 5-minute intervals. The nuclei were counterstained with DAPI, and photos were taken by Olympus IX71 microscope equipped with DP72 digital camera.

### 2.5. Enzyme Linked Immunosorbent Assay (ELISA)

To analyze the amount of cytokines secreted in the culture media, supernatants from the cells were collected during differentiation at the distinct days (D0, D8, D14, and D25) and determined by ELISA for interleukin-6 (IL-6; Platinum ELISA, eBioscience, Austria) and interleukin-8 (IL-8; Platinum ELISA, eBioscience, Austria) according to the manufacturer’s instructions.

### 2.6. Statistical Analysis

All quantitative experiments, including qRT-PCR and ELISA, were performed using 10 biologically independent replicates. Significant differences between groups were examined by analysis of variance (ANOVA) and the Student’s t-test. Differences were considered to be statistically significant at P < 0.05. Data were presented as mean±SD. Pearson’s correlation was used to investigate the relationship of gene expression between TLR2 and TLR4 during differentiation.

## 3. Results

### 3.1. TLRs expression in ESCs derived cardiomyocyte

The mRNA expression of TLRs in ESC-derived cardiomyocytes was examined by RT-PCR. Specific primers were used to amplify sequences of TLRs, and the identity of the fragments was confirmed by sequencing. According to gel electrophoresis in Figure 1, the results showed that all of the TLRs were present in ESC-derived cardiomyocytes. GAPDH has been considered as an internal control in this study (Fig. 1A).

**Figure 1.**
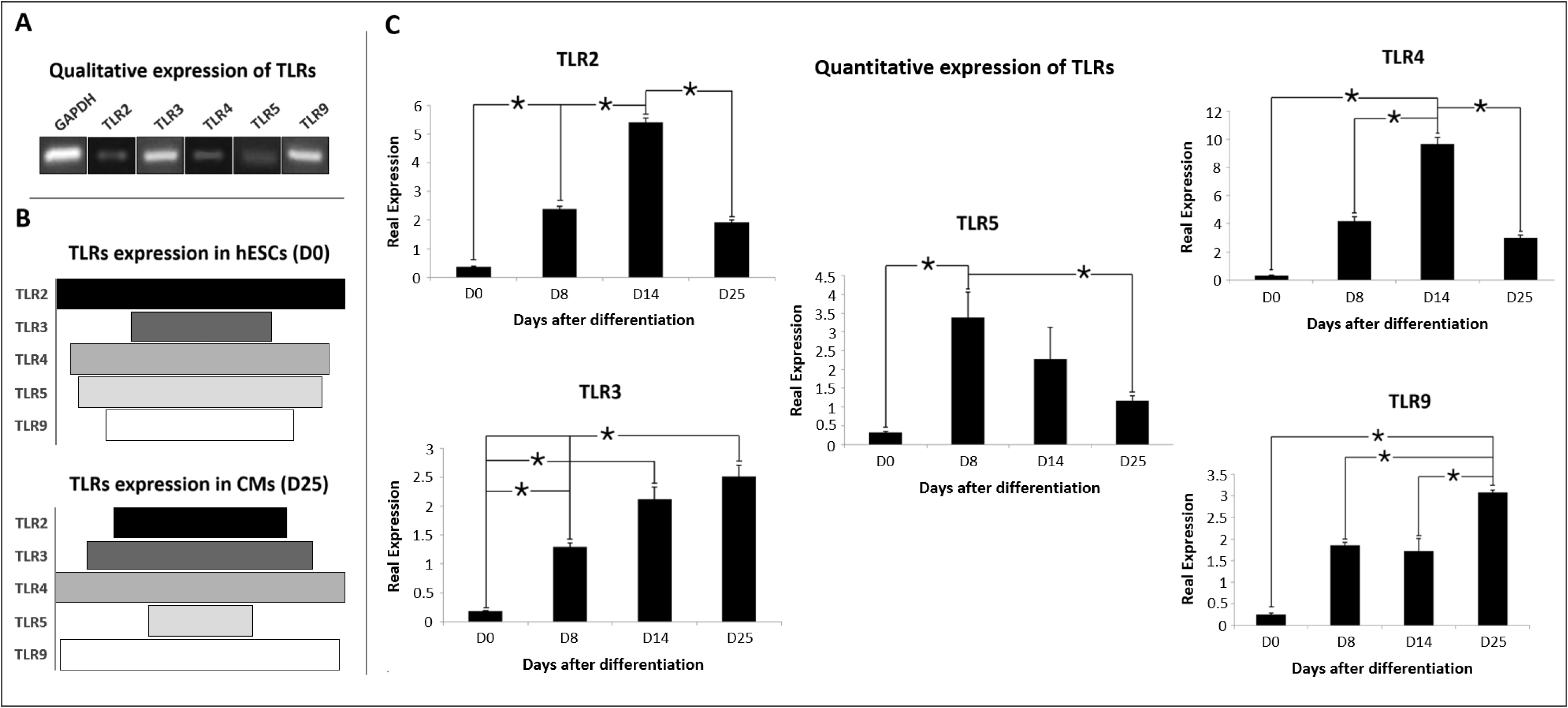
Qualitative expression of TLRs in ESC-derived cardiomyocytes and quantitative expression of TLRs during cardiomyocyte differentiation. (A) After 35 cycles of PCR and gel electrophoresis of the PCR product, the results showed that the mRNAs of TLR2, TLR3, TLR4, TLR5, and TLR9 were expressed in cardiomyocytes differentiated from ESCs. GAPDH has been considered as an internal control. (B) A quantitative expression comparison of various TLRs on the day null of differentiation (D0) in hESCs, and on the last day of differentiation (D25) in fully differentiated cardiomyocytes. (C) Quantitative expression of cell membrane TLRs (TLR2, TLR4, and TLR5) and endosomal TLRs (TLR3 and TLR9) during differentiation. The experiment was performed for at least 10 independent biological replicates. *P<0.05. Data are shown as mean±SD.

### 3.2. TLRs expression was varied during cardiomyocyte differentiation

The process of TLRs mRNA expression during cardiomyocyte differentiation was evaluated by quantitative RT-PCR. According to Q-PCR data, the expression pattern of TLR2 and TLR4 was the same (Fig. 1C), and they showed the strongest correlation with each other among TLRs (Fig. 2A, B). Moreover, both represented the lowest expression in mature cardiomyocytes and the highest expression on day 14 during differentiation (Fig. 1C). TLR5 expression decreased from day 8 to day 25. In contrast, TLR3 expression increased during the differentiation process. (Fig. 1C). TLR9 expression showed the highest expression in day 25 (mature cardiomyocyte) (Fig.1C), and it represented close correlation in gene expression with TLR3 (Fig. 2A). TLRs expression showed that TLR4 has the highest expression and TLR3 has the lowest expression during cardiomyocyte differentiation (Fig. 3A). Furthermore, Comparison of TLRs expression in different days indicated that the day 14 is a critical time during differentiation with the highest expression of TLRs (Fig. 3B).

**Figure 2.**
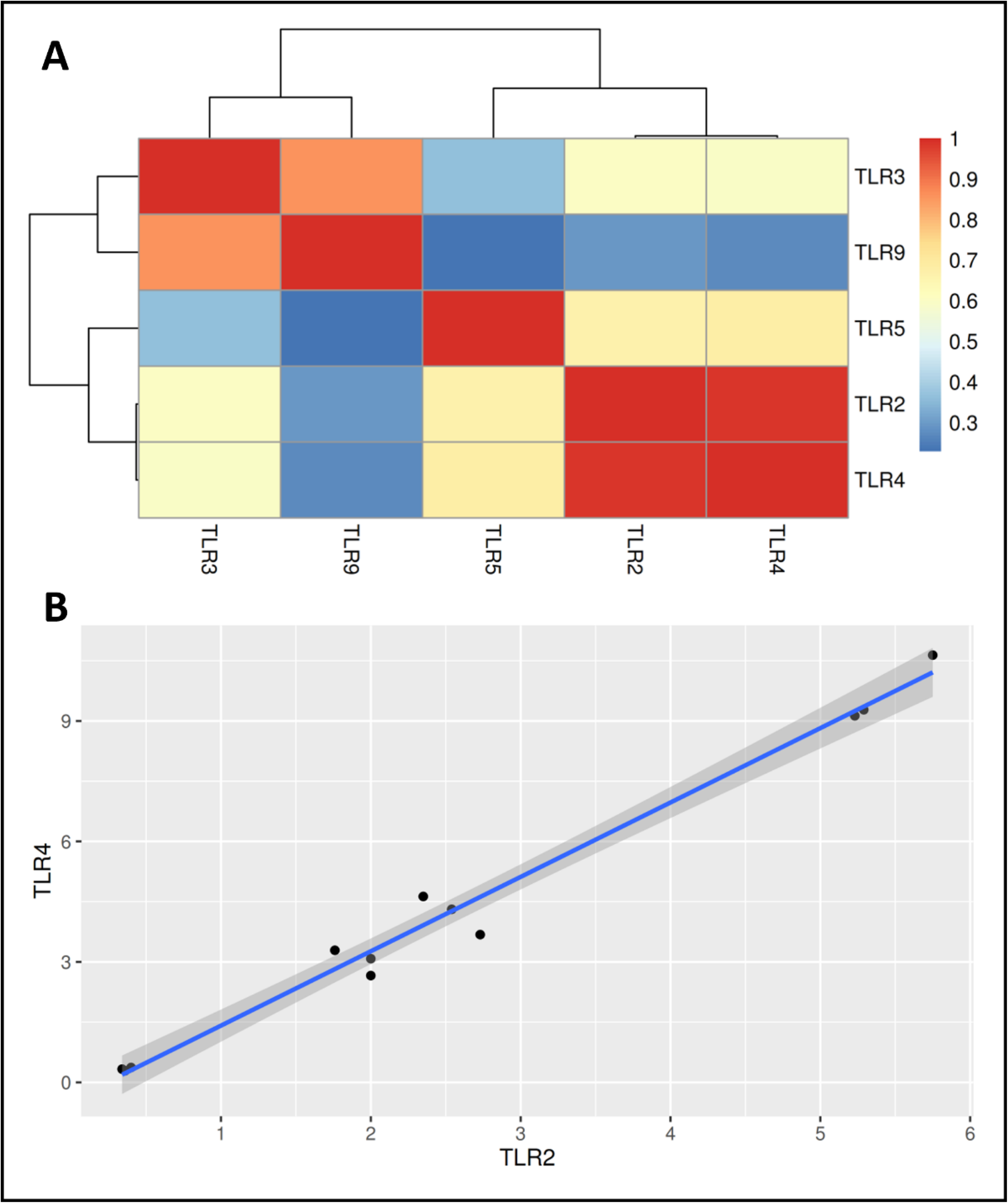
Correlation between TLR expression in CM differentiation. **(A)** TLR2 and TLR4 show the strongest correlation. After these TLRs, TLR3 and TLR9, as endosomal receptors, exhibit the closest correlation in gene expression. (Red color shows the highest, and the blue color represents the lowest correlation in the heatmap. **(B)** Correlation of gene expression between TLR2 and TLR4. Pearson correlation coefficient (r) is +0.94 between these two receptors.

**Figure 3.**
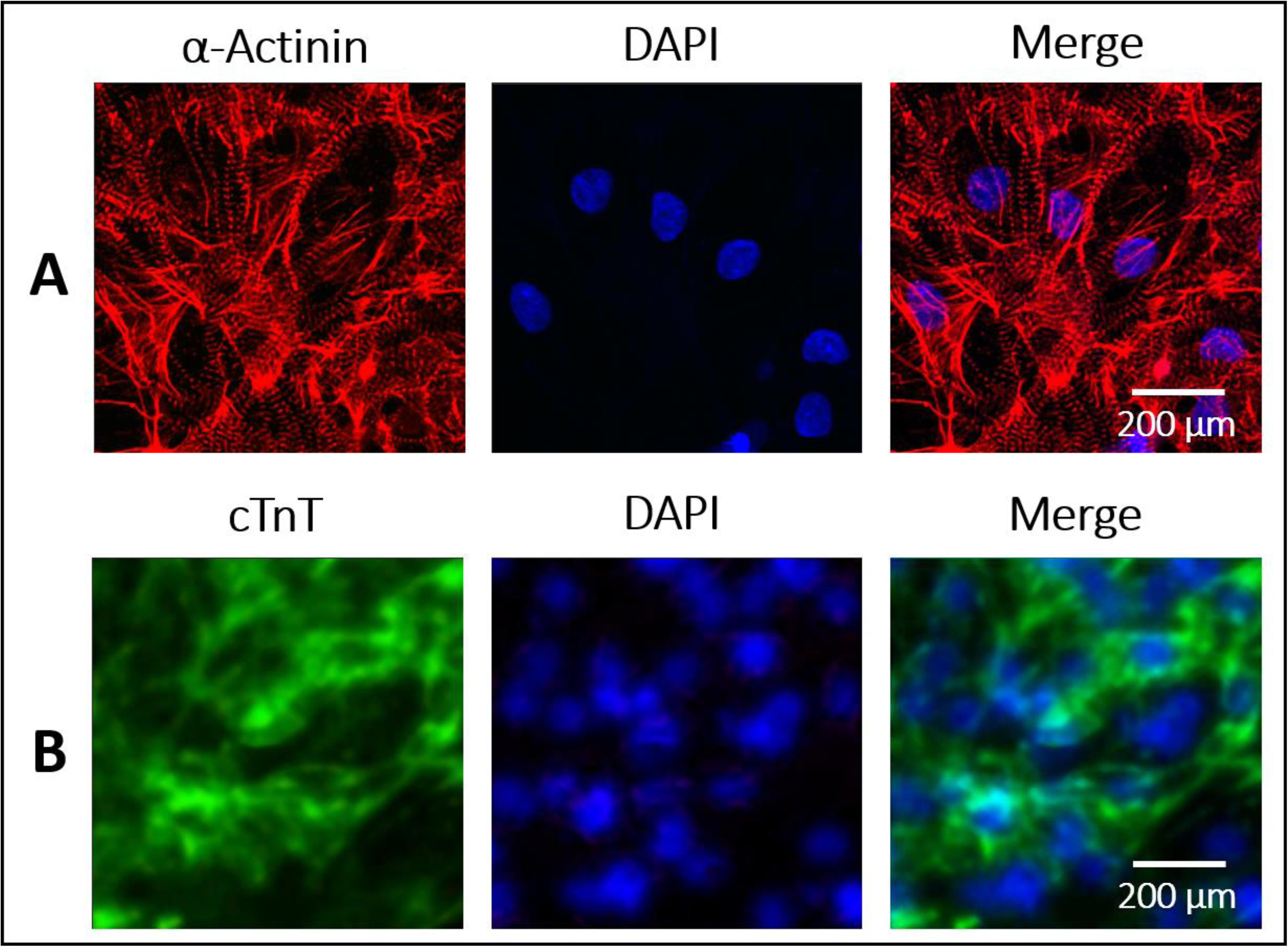
Marker analysis of hESC-CM. Immunostaining of plated hESC-CMs at day 25 of differentiation for cardiomyocyte-specific markers showed **(A)** α-actinin (Red) magnification (10X), and **(B)** cTnT (Green) magnification (20X). Nuclei were counterstained with DAPI (Blue).

### 3.3. Confirmation of CM differentiation by expressing cytoskeletal proteins

Immunostaining data showed that emryonic stem cells derived cardiomyocytes express cytoskeletal proteins such as alpha actinin (α-actinin) (Fig. 4A) and cardiac troponin T (cTnT) (Fig. 4B). The expression of these proteins exhibits the striation pattern in differentiated cardiomyocytes.

**Figure 4.**
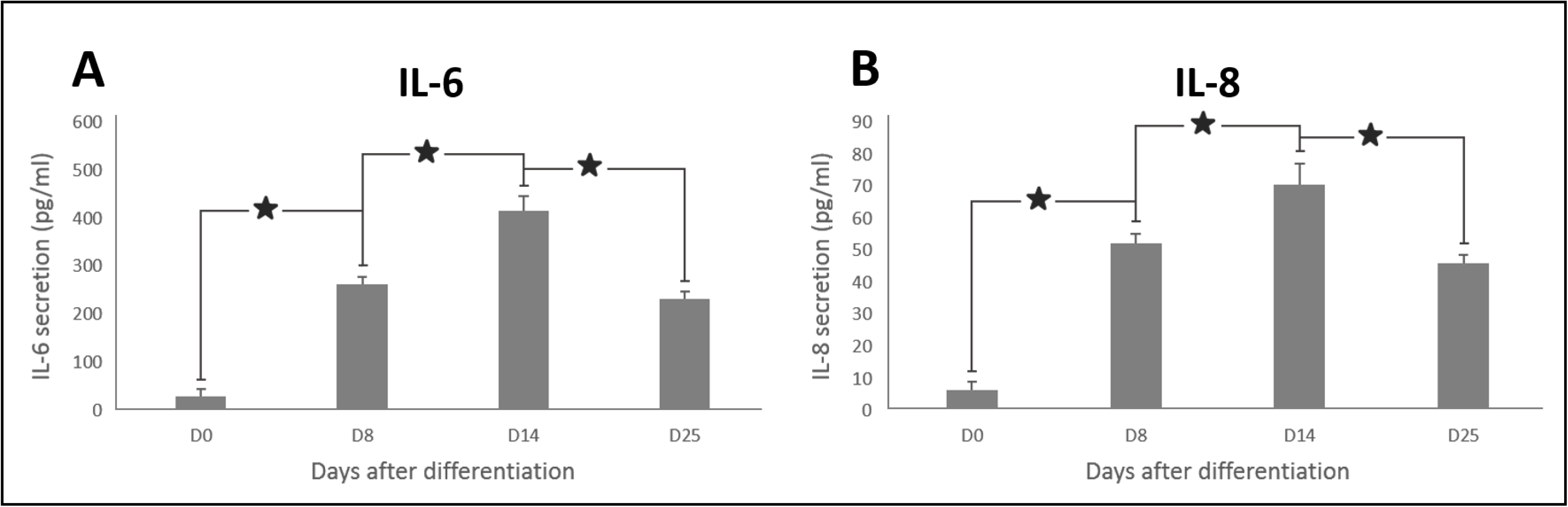
IL-6 and IL-8 expression during differentiation. ELISA test for assaying the secretion of two main important cytokines **(A)** IL-6, and **(B)** IL-8. *P<0.05. Data are reported as mean±SD.

### 3.4. IL-6 and IL-8 secretion during differentiation

The expression of two main proinflammatory cytokines secreted after induction of both cell membrane and endosomal TLRs, IL-6 and IL-8, was determined by ELISA. Both cytokines showed a pattern like TLR2 and TLR4 expression during differentiation (Fig. 5A, B). It is clear that the maximum secretion occurred on day 14 of cardiomyocyte differentiation and their expression continued until the cells were fully differentiated into cardiomyocytes.

## 4. Discussion

Treatment of cardiac diseases with hESC-derived cardiomyocytes is now close to reality; however, little is known of their responses to physiological and pathological insult. To explore such responses, understanding the immune response during cardiomyocyte differentiation from hESCs is quite indispensable. Innate immunity and inflammatory responses have been implicated in myocardial ischemia/reperfusion (I/R) injury and other heart diseases [26]. As a rule of thumb, the expression of TLRs as main receptors in innate immunity and the link between innate and adaptive immunity, stimulates cell inflammation and cardiac remodeling [7, 13] and promotes injury in the heart in response to ischemia, ischemia-reperfusion, or hypertrophic stimuli [27]. They are expressed in the cardiovascular system and could thus be a key link between CVDs and the activation of the immune system [28, 29]. Not only did we pursue the evaluation and comparison of different TLR expression during CM differentiation from human ESCs as a goal, but obtaining reliable cardiomyocytes regarding immunological parameters and responses that can be used in regenerative therapy was also an important goal for us in this study. To do this, TLR2, TLR4, and TLR5 were representative of plasma membrane receptors, and TLR3 and TLR9 were studied as endosomal TLRs. Based on the results, the expression pattern of TLR2 and TLR4 showed a strong correlation, while TLR3 and TLR9 represented the same pattern of expression during differentiation.

According to other studies, some of the TLRs are expressed in undifferentiated hESCs. Foeldes and his colleagues demonstrate that TLR1, TLR3, TLR4, TLR5, and TLR6 were expressed in hESCs [30]. Our data confirmed these results for TLR3 to TLR5. Moreover, the expression of TLR2 and TLR9 in these cells was reported in our study. TLRs are also readily detectable in cardiac myocytes [31]. According to Feng and Chao’s study, TLR2, TLR3, TLR4, TLR5, TLR7, and TLR9 are expressed in mouse heart tissue and in the CM cell line [32]. Another study reported that TLR2, TLR3, TLR4, TLR6, TLR7 and TLR9 are expressed in ventricular myocytes, whereas TLR1 and TLR5 are not [5]. Our data confirm these results regarding the expression of TLR2, TLR3, TLR4, and TLR9. In contrast, we depicted TLR5 expression in CMs.

The study of TLR expression during differentiation is very interesting for therapeutic targets. The TLR pathway is involved in the development of the central nervous system [33]. Additionally, the role of these receptors during the differentiation of human aortic endothelial cells from ESCs has been evaluated [30]. In this study, the expression pattern of TLRs during differentiation of CMs from hESCs at the mRNA level was reported for the first time. In comparison to Hsueh’s study, we can categorize our differentiation process into four distinct stages, in which the day zero is a pluripotent state, the germ layer specification is equal to the 8th day, the appearance of progenitors occurs around day 14 when the commitment is started, and lastly, mature CMs emerge on the 25th day [34]. Except for the Pluripotent state, where TLR4 was less expressed than TLR2, in all subsequent stages, TLR4 represented a higher level of expression than the other TLRs evaluated in our study. It is likely due to the more widespread signaling pattern of TLR4 than the other TLRs. By interacting with CD14, a co-receptor of many TLRs, transferrin silences the TLR signaling complex, especially TLR4 [35]. Moreover, Zhang and colleagues reported that transferrin improves the CM differentiation from human pluripotent stem cells [36]. Regarding these two roles of transferrin, it is inferred that high expression of TLR4 is not encouraging of CM differentiation. Interestingly, our results corroborated these data, where TLR4 was upregulated until the initiation of the commitment stage at day 14; however, its expression decreased in the subsequent days of differentiation and hit its lowest level in the mature cardiomyocyte.

After coronary artery ligation, mortality and left ventricular dilatation were significantly reduced, and left ventricular function was preserved in TLR2 _/_ mice compared with wild type mice. Furthermore, left ventricular remodeling and survival are improved in TLR4-deficient mice, accompanied by a reduction in pro-inflammatory cytokines and alterations in extracellular matrix remodeling [37, 38]. These evidences prove that inhibition of TLR2 and TLR4 signaling could reduce cardiac damage after ischemic injury. Accordingly, it is expected that the expression of TLR2 and TLR4 must decrease during cardiomyocyte differentiation. Our data not only confirmed the same expression pattern of TLR2 and TLR4, but also illustrated a decrease in their gene expression between the third and last stages of differentiation. Also, Boyd and his colleagues showed that ligand activation of TLR2, TLR4, and TLR5, but not TLR3, TLR7, or TLR9, resulted in cardiomyocyte expression of the inflammatory cytokine interleukin 6 (IL-6), the chemokines KC and macrophage-inflammatory protein-2 (MIP-2), and the intercellular Adhesion Molecule 1 (ICAM1). Activation of these Toll-like receptors via NF-κB signaling was associated with decreased cardiomyocyte contractility [5]. The same expression pattern of IL-6 and IL-8 with TLR2 and TLR4 during CM differentiation in our study confirms Boyd’s results. Moreover, both mentioned ILs have exhibited their minimum secretion during differentiation in fully differentiated cardiomyocytes, as indicated by the minimum level of expression of TLR2, TLR4, and TLR5 in the last stage of differentiation.

Cardiomyocyte maturation is a critical point that has recently drawn the attention of scientists due to the maturation defects in PSC-CM and its potential role in cardiac disease [39]. To improve the maturation of PSC-CMs, various methods, including 3-dimensional structures, mechanical stimuli, genetic regulation, and chemical agents, have been recruited [34, 40]. According to Hodgkinson’s study, the blocking of CM maturation, specifically the failure of maturation gene expression in precursor cells committed to the CM line, was found upon TLR3 inhibition [41]. This positive correlation between the expression of TLR3 and CM maturation is confirmed by our results, where the expression of this receptor represented an upward trend during differentiation and reached its highest expression in the fully differentiated CM. This suggests that the yielded CMs in our study are mature.

TLR9, in immune cells, induces inflammation upon tissue injury, whereas this receptor plays a key role in energy metabolism and cellular protection in non-immune cells such as CMs and neurons. TLR9 stimulation reduced energy substrates and increased the AMP/ATP ratio, subsequently activating AMP-activated kinase (AMPK), leading to increased stress tolerance against hypoxia in cardiomyocytes without inducing the canonical inflammatory response [42]. Accordingly, TLR9 upregulation during cardiomyocyte differentiation and reaching the highest expression in fully differentiated CMs is expected, as our data strongly corroborates. As key receptors with widespread roles in innate immunity and initiating adaptive immunity, TLRs have footprints in cardiovascular ischemia diseases, cellular behaviors, cardiomyocyte protection, contractility, and even maturation. According to recent literature, the role of TLRs in cellular differentiation has been reported; however, there is no evidence evaluating the expression of these critical receptors during cardiomyocyte differentiation, which is considered an innovative aim of this study. Decreasing expression of cell membrane TLRs (TLR2 and TLR4) after generating progenitors around day 14 until fully differentiated CMs, besides high expression of endosomal TLRs (TLR3 and TLR9) in hESC-CMs, are positive clues, each of which represents a characteristic of fully functional differentiated cardiomyocytes, which could be a golden choice to apply in regenerative therapy. Hence, deciphering the expression pattern of TLRs during cardiomyocyte differentiation can pave the way for obtaining optimized functional cardiomyocytes and be introduced as a novel procedure for CM characterization, associated with other prevalent methods of characterization. Moreover, to obtain highly functional CMs, TLR expression optimization is highly recommended. For instance, as cardiomyocyte maturity is the missing link in all current differentiation methods, the upregulation of TLR3 likely helps us to generate highly mature CMs.

## 5. Acknowledgments

All authors have contributed actively to prepare this article and their roles are as follows: Mohammad Reza Hashemzadeh: conception, design, acquisition of data, analysis and interpretation of data, drafting the manuscript / Samane Aghajanpour: acquisition of data, analysis and interpretation of data / Nasser Aghdami: drafting and revising the manuscript / Reza Aflatoonian: design, analysis and interpretation of data, drafting and revising the manuscript.

